# Elucidating dispersal ecology of reclusive species through genetic analyses of parentage and relatedness: the island night lizard (*Xantusia riversiana*) as a case study

**DOI:** 10.1101/125666

**Authors:** Stephen E Rice, Rulon W Clark

## Abstract

Characterizing dispersal and movement patterns are vital to understanding the evolutionary ecology of species. For many reclusive species, such as reptiles, the observation of direct dispersal may be difficult or intractable. However, dispersal distances and patterns may be characterized through indirect genetic methods. We used genetic and capture data from the island night lizard (*Xantusia riversiana*) to estimate natal dispersal distances through indirect genetic methods, characterize movement and space use patterns, and compare these distances to previous estimates made from more traditional ecological approaches. We found that indirect estimates of natal dispersal were greater than previous field-based estimates of individual displacement of 3-5 m. Parent-offspring differences had a mean of approximately 14 m on Santa Barbara Island (SBI) and 41 m on San Clemente Island (SCI) whereas Wright’s σ was estimated at 16 m on SBI and 20 m for SCI. Spatial autocorrelation with correlograms of Moran’s I revealed large differences in the scale of autocorrelation between islands (SBI=375 m, SCI=1,813 m). Interpretation of these distances as average per generation distance of gene flow was incongruent with parentage analyses and σ. We also used variograms to evaluate the range of spatial autocorrelation among two inter-individual genetic differences. The range of spatial autocorrelation again identified different scales on the two islands (102 - 169 m on SBI and 955 - 1,424 m on SCI). No evidence of sex-biased dispersal was found on either island. However, a permutation logistic regression revealed that related individuals >0.8 years old were more likely to be captured together on both islands. Overall, our findings suggest that field-based estimates of individual displacement within this species may underestimate genetic dispersal. We suggest indirect inferences of natal dispersal distances should focus on parentage analyses and Wright’s σ for parameter estimation of individual movement, whereas the ranges identified by spatial autocorrelation and variograms are likely to be relevant at the metapopulation or patch scales. Furthermore, characterization of capture patterns and relatedness revealed kin-affiliative behavior in *X. riversiana*, which may be indicative of delayed dispersal and cryptic sociality. These results highlight the power of parentage- and relatedness-based analyses for characterizing aspects of the movement ecology of reclusive species that may be difficult to observe directly. These data can then be leveraged to support future conservation and population modeling efforts and assess extinction risks and management strategies.

## INTRODUCTION

The dispersal and movement of organisms is a fundamental process in population biology, yet may be difficult to characterize even in abundant species. Dispersal studies provide insight into the evolutionary ecology of focal species and inform conservation planning and management (e.g. Bowler & Benton 2005; Dussex et al. 2016; Hawkes 2009). Dispersal, defined here as the movement of individuals away from their natal sites, may be quantified directly through long-term capture-mark-recapture (CMR) studies and spatial monitoring, or indirectly through genetic inference methods.

Both approaches to characterizing dispersal have limitations. Direct observations of dispersal are labor intensive and time consuming, resulting in reduced sample sizes that may underestimate typical dispersal rates and distances (e.g. Dussex et al. 2016), whereas genetic inference may be complicated through modeling assumptions and challenges in study design and sample collection (Broquet & Petit 2009). While few studies use both direct and indirect inference methods, the proliferation of genetic methods has led to a variety of analytical approaches for detecting dispersed individuals (Manel et al. 2005), categorize age and sex biases in dispersal patterns, and characterize dispersal distances (Broquet & Petit 2009; Goudet et al. 2002). The processes and potential confounds of these methods are important to understand, as incorporating unrealistic assumptions or parameters within spatial models of dispersal may yield inaccurate results and directly impact management actions and efficacy (Bowler & Benton 2005; Hawkes 2009). However, for reclusive species the characterization of dispersal from field data may be especially problematic due to low probabilities of recapture (e.g. Pimm et al. 2015); thus indirect methods present a compelling tool for characterizing dispersal patterns and distances.

Genetic methods are an increasingly common tool for elucidating species dispersal ecology. Recently, Moore et al (2014) used parentage analyses with pairwise distances between dyad members to characterize condition-dependent dispersal patterns in American black bears. Dussex et al (2016) found that assignment methods were generally inconsistent with CMR and parentage analyses inferences. Furthermore, they found that parentage analyses were the most powerful approach at fine scales for elucidating dispersal ecology of the greater white-toothed shrew. In addition to understanding dispersal, managers often need to determine the spatial extent over which landscape influences a focal species (Jackson & Fahrig 2014). Jackson and Fahrig (2014) found that the scale at which landscape structure affects species varies with the population outcome measured. The simulation study conducted by Jackson and Fahrig (2014) suggested that the scale of landscape structure for population persistence should be a lower bound for conservation as the scale needed for supporting genetic diversity is much larger. Taken together, statistical methods that help characterize the dispersal of species and the scale at which landscape affects genetic structure can provide vital information in the context of conservation.

The island night lizard (*Xantusia riversiana*) was recently delisted from the Endangered Species Act and provides a unique opportunity to evaluate the utility of indirect methods to elucidate dispersal ecology. This species has been well studied, has few documented predators, and exists in discrete insular populations with high regional abundances of 3,200 individuals/ha in prime habitat. Even with long-term study, direct observations of individual movement suggest very small individual displacement distances of 3 to 6 m over multiple years (Fellers & Drost 1991; Mautz 1993). The dearth of information on island night lizard dispersal ecology is a direct obstacle to modelling metapopulation dynamics and potential threats due to climatic change.

We applied genetic inference methods to characterize dispersal ecology and infer natal dispersal distances in *X. r. reticulata*. We used genetic and capture data from our landscape genetics analysis (Rice & Clark 2016) to characterize dispersal ecology and space use on two of the three California Channel Islands this species is known to occupy. The goal of this study was to leverage existing data to assess different techniques for estimating dispersal from genetic data and compare these patterns to values derived from ecological studies.

## METHODS

### Study System and Data

Island night lizards are endemic to three California Channel Islands, two of which were evaluated by Rice and Clark (2016): Santa Barbara Island (SBI) and San Clemente Island (SCI). In brief, we captured 917 island night lizards across both islands utilizing a clustered sampling approach. Individuals were genotyped at 23 microsatellite loci (Rice et al 2016) and first-order relatives, defined as parent-offspring and full siblings, were identified using a consensus approach between three methods: COLONY vers 2.0.5.9 (Jones & Wang 2010), CERVUS vers 3.07 (Kalinowsi et al 2007), and the DyadML estimator (Milligan 2003) as calculated in COANCESTRY vers 1.0.1.5 (Wang 2011). The current study draws on the capture data, individual genetic profiles, relatedness and relationship analyses of Rice and Clark (2016).

### Statistical Approaches

We used 4 approaches to quantify dispersal distances in the island night lizard: pairwise distances between inferred relationships in Rice and Clark (2016), estimation of Wright’s gene-dispersal distance (Wright 1946), correlogram of Moran’s I (reviewed in Hardy & Vekemans 1999), and range estimates from variograms (reviewed in Wagner et al. 2005). We characterized sex-biases in dispersal using the approach of Goudet et al. (2002). We evaluated predictors of co-capture among individuals with a permutation based logistic regression on distance matrices (LRDM, Prunier et al. 2015). Statistical analyses were carried out in R (R Core Team 2016) using the packages *adegenet* (Jombart 2008), *hierfstat* (Goudet & Jombart 2015), *coin* (Hothorn et al. 2006), *phylin* (Tarroso et al. 2015), and *fmsb* (Nakazawa 2015). We estimated Wright’s gene dispersal distance, σ, and Moran’s I using the program SPAGEDI vers 1.5 (Hardy & Vekemans 2002).

### Distances Between First-Order Relatives

The pairwise distances between parents and offspring have been demonstrated to be a powerful tool in the characterization of dispersal ecology (e.g. Dussex et al. 2016; Moore et al. 2014). We utilized the data of Rice and Clark (2016) to characterize distances between inferred relationship classes. First-order relatives consisted of individuals identified through the consensus method in Rice and Clark (2016) wherein dyads were considered first-order relatives when the inferred relationship was parent-offspring, full sibling, or the relatedness coefficient, DyadML, was greater than 0.35. Relationship classes of parent-offspring, full sibling, and half sibling were inferred by COLONY. Unrelated samples were all relationships not detected by COLONY.

We used an approximate general independent test of distances between ordered relationship groups in the package *coin* to test whether pairwise geographic distances differed between relationship groups within and between islands. To correct for the presence of neonates captured in close proximity to adults, we present analyses based on the full data sets and data sets consisting only of individuals greater than 40 mm snout-to-vent length (SVL) which equates to approximately 0.8 years old (Fellers & Drost 1991).

### Wright’s σ

Wright (1946) described an isolation by distance model in which the genetic neighborhood is a 2-dimensional area in which most mating events occur. This model can be used to estimate σ from a regression of the inter-individual genetic distance, Rousset’s a (Rousset 2000), on the logarithm of the inter-individual distance (Hardy et al. 2006; Rousset 2000). We assumed drift-migration equilibrium and estimated σ, interpreted as mean natal dispersal distance, for each island across 3 distance classes and 5 density estimates (Table 1). Density estimates were based on the average effective population size per sample site as estimated by the linkage-disequilibrium method (Hill 1981) and confidence intervals from the program NeEstimator vers 2.01 (Do et al. 2014). Additional estimates of density were derived from the census population size over the entire area of each island and the prime-habitat area listed in the United States Fish and Wildlife Service post-delistment monitoring plan (USFWS 2014).

**Table 1:**
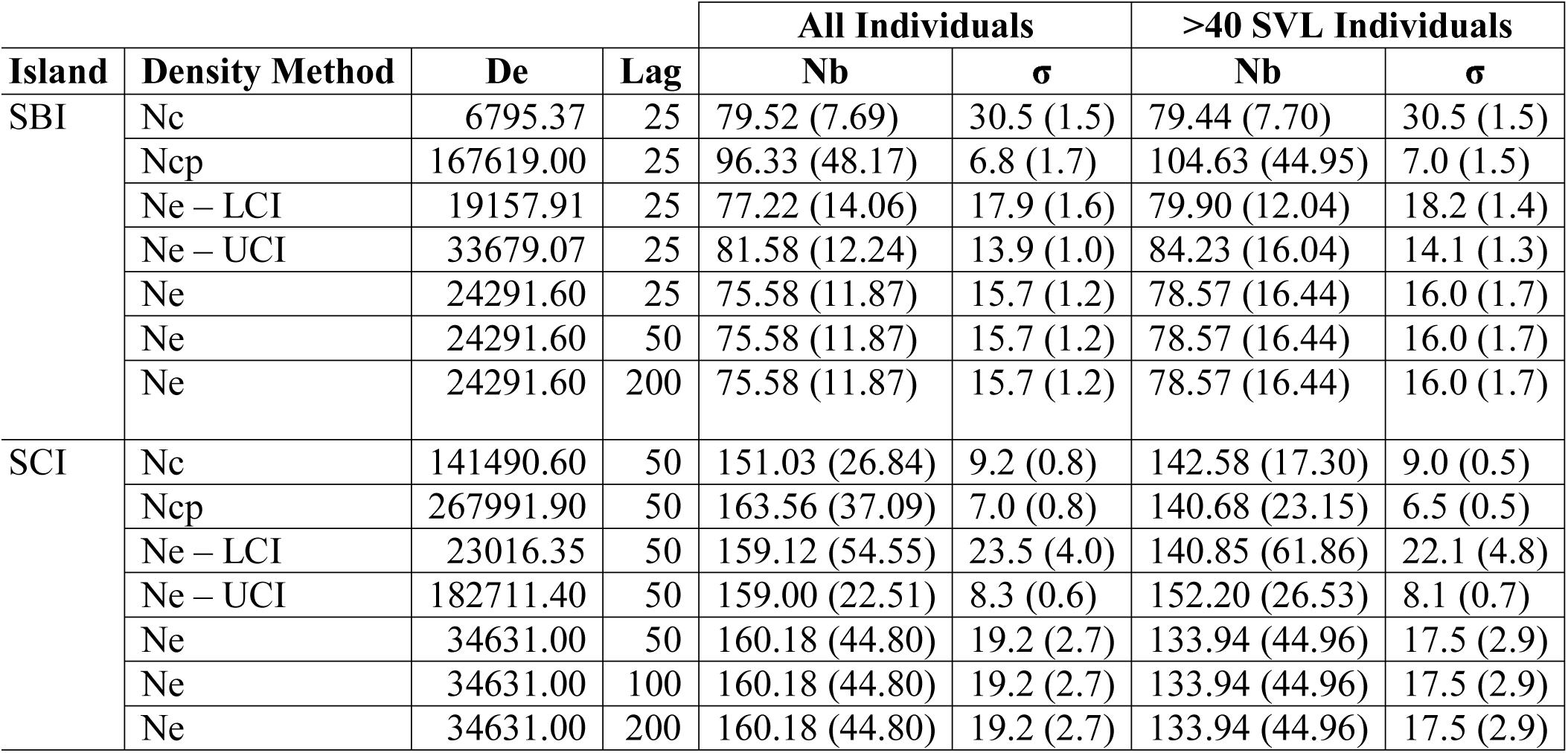
Isolation by Distance Model Results. Results from the program SPAGEDI for estimation of Wright’s σ. Islands were estimated independently (SBI=Santa Barbara Island, SCI=San Clemente Island). Density (De) was estimated as individuals/km^2^ by considering the census population size (Nc) and island area, average for effective density across sampling locations (Ne) with 95% confidence intervals (Ne-LCI=lower, Ne-UCI=upper), and census population size over prime habitat area (Ncp). Lag distance refers to the increment of distance bins (m). SPAGEDI estimated the number of breeders (Nb) and σ (m); standard errors in parentheses for each parameter were estimated by jackknifing over loci. Two data sets were ran for each island, the full data set (All Individuals) and a data set in which the young of year were removed (>40 SVL Individuals).

### Moran’s I

Moran’s I is a common measure of spatial autocorrelation of individual allele frequencies (Hardy & Vekemans 1999). Moran’s I was calculated for each pairwise distance bin, set at increments of 150 m for SBI and 500 m for SCI, and significance was tested with 1000 permutations. The increments for distance bins were assigned based on the smallest distance at which all bins had observations. For each island, we produced a correlogram of Moran’s I at each distance class. The distance at which the value of Moran’s I became ≤ 0 was interpreted as the point at which individual allele frequencies were no longer spatially autocorrelated. We interpreted this point as the average maximum dispersal distance of island night lizards (e.g. Yaegashi et al. 2014).

### Variograms

Variograms assess the spatial autocorrelation of a variable by depicting the semivariance (defined as half the variance of all pairwise differences) against distance, and may identify the spatial scale of dispersal processes (Dutech et al. 2008; Le Corre et al 1998). The empirical data is used to generate a variogram which is then fit with a theoretical variogram with parameters for nugget (semivariance associated with non-spatial effects), sill (the value at which semivariance stabilizes), and range (the scale of effect or threshold of spatial independence) (Wagner et al. 2005). There are few guidelines for determining the increment or maximum distances considered in empirical variograms; therefore, we followed the conventions of using the minimum lag distances which produced a minimum of 30 observations per bin and limited variograms to one-half the maximum pairwise distance compared (Rossi et al. 1992). Lag distances differed between Moran’s I and variogram analyses, due to variogram constraint to a smaller maximum distance.

We used the package *phylin* to produce empirical and theoretical variograms. For theoretical variograms, exponential models were fit to all data sets with the nugget set to the semivariance of the first distance bin and the remaining parameters estimated within the package. The variogram approach implemented in *phylin* requires pairwise distance variables; therefore we evaluated DPS, an inter-individual genetic distance calculated as 1 – proportion of shared alleles in *adegenet*, and a measure of (un)relatedness by subtracting DyadML values from 1, the theoretical maximum probability under identity by descent (Milligan 2003).

### Sex-Biased Dispersal

The methods of Goudet et al (2002) identify a permutation t-test approach to test for sex-biases in dispersal based on four metrics derived from genetic data of each sex: mean and variance of the corrected assignment index (mAIc and vAIc, respectively), Fst, and Fis.

However, these methods have been reported to perform well only in the presence of strong sex-biased dispersal (Goudet et al. 2002). Statistical tests to detect sex bias were conducted in the package *hierfstat* with the metrics mAIc, vAIc, Fst, and Fis (Goudet et al. 2002). Significance for comparisons based on mAIc and vAIc were ran with 10,000 permutations whereas tests based on Fst and Fis were based on 1,000 permutations due to computational constraints.

### LRDM

We used an extension of permutation-based LRDM (Prunier et al. 2015) to assess whether the probability of capturing individuals together could be attributed to the predictors of pairwise sex, sexual maturity, relatedness, or relationship. We used a binary response variable with success defined as being captured together. Predictors were pairwise distance matrices with sex and sexual maturity coded as categorical comparisons.

Models were constructed from single variable up to full models; due to the highly collinear nature of DyadML and COLONY -inferred relationships (data not shown) these predictors were not included in the same models. Logistic regression models were constructed with the glm function and a binomial ‘logit’ link. Nagelkerke’s R^2^ (Prunier et al. 2015; Smith & McKenna 2013) was calculated for each model using the NagelkerkeR2 function in the package *fmsb* and served as the reference distribution statistic for determining significance by permuting the binary response matrix 10,000 times and recalculating Nagelkekre’s R^2^ for each permutation. Semi-standardized beta weights for the full models were calculated as in Prunier et al. (2015) with the odd’s ratios calculated as the exponentiated semi-standardized beta weights following King (2007).

## RESULTS

Mean pairwise distances (Table 2) were significantly different for each relationship group within both islands (p < 2.2 × 10^-16^). Mean pairwise distances for full siblings and parent offspring were not significantly different within either island (SBI p=0.1017, SCI p=0.7566); all other comparisons were significant at the p<0.005 level. When controlling for young of year (SVL < 40 mm), mean pairwise distances were significantly different between islands for first-order relationship (p=0.0062) and parent-offspring dyads (p=0.0228). When young of year were included there were no significant differences. When controlling for young of year, mean parent-offspring distance for SBI was 13.93 m and 40.68 m on SCI.

**Table 2:**
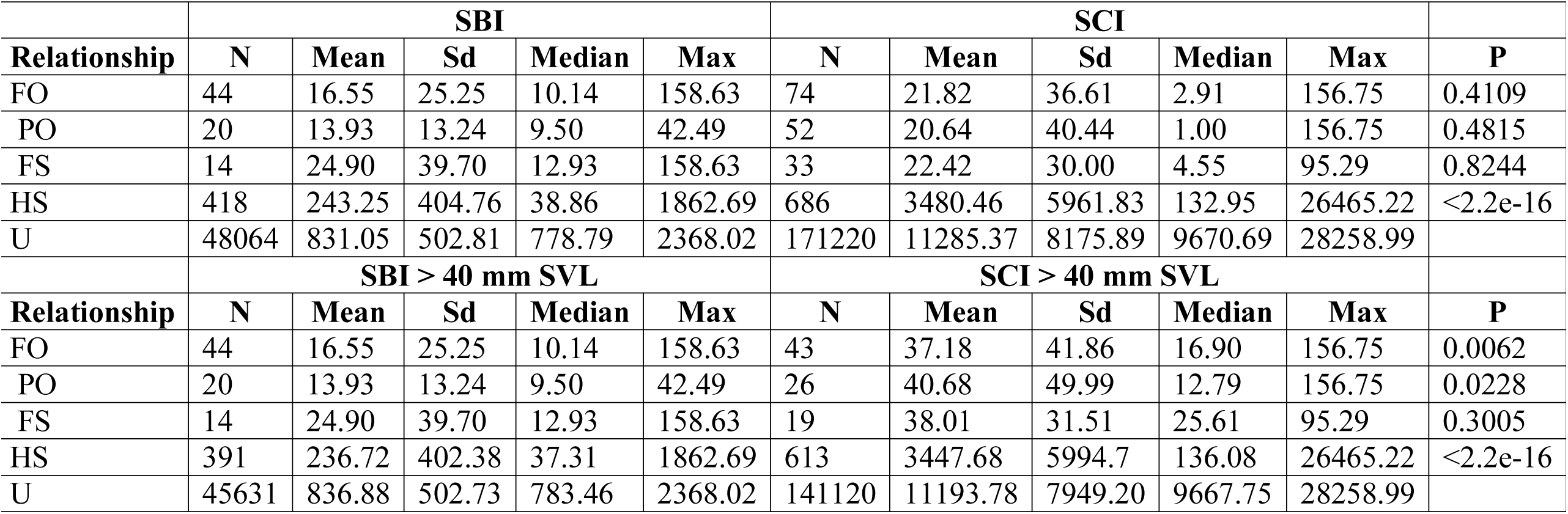
Pairwise Distances among Relatives. Islands are indicated as Santa Barbara Island (SBI) or San Clemente Island (SCI). Relationship were first-order (FO, based on Rice and Clark (2016)), COLONY inferred relationships of parent-offspring (PO), full sibling (FS), half sibling (HS) or unrelated (U). For each relationship, the number of pairwise comparisons (N), mean (Mean), standard deviation (Sd), median (Median), and maximum (Max) distance values (m). The P column is the approximated p-value comparing the same Relationships between islands. The lower half of the table displays values for each island with young of year samples (>40mm SVL) removed.

The gene dispersal estimate, σ, ranged between approximately 7 m and 31 m for SBI and approximately 7 m to 23 m for SCI. Estimates of σ were sensitive to density estimates. However, estimates of σ were consistent across different distance classes when density was constant and robust to the inclusion of young of year (Table 1). Constraining density estimates to average effective density (Ne/km^2^) across sample locations resulted in estimates of 16 m for SBI and 19 m for SCI for full data sets and 16 m for SBI and 18 m for SCI when young of year were removed.

Spatial autocorrelation on SBI, as represented by Moran I, was significantly positive (p≤0.005) at distances less than 375 m at which Moran’s I became negative but not significant (Figure 1). On SCI, spatial autocorrelation was significantly positive (p≤0.001) up to distances of 1,395 m and had standard errors overlapping 0 at a distance of 1,813 m (Figure 2). Variograms based on relatedness had higher support, as indicated by R^2^, than those based on DPS for both islands (Figures 3,4). On SBI, the range for the DyadML variogram was approximately 102 m whereas SCI had a range estimate of 955 m. The genetic distance measure, DPS, resulted in greater range estimates on SBI at 169 m and SCI at 1424 m.

**Figure 1:**
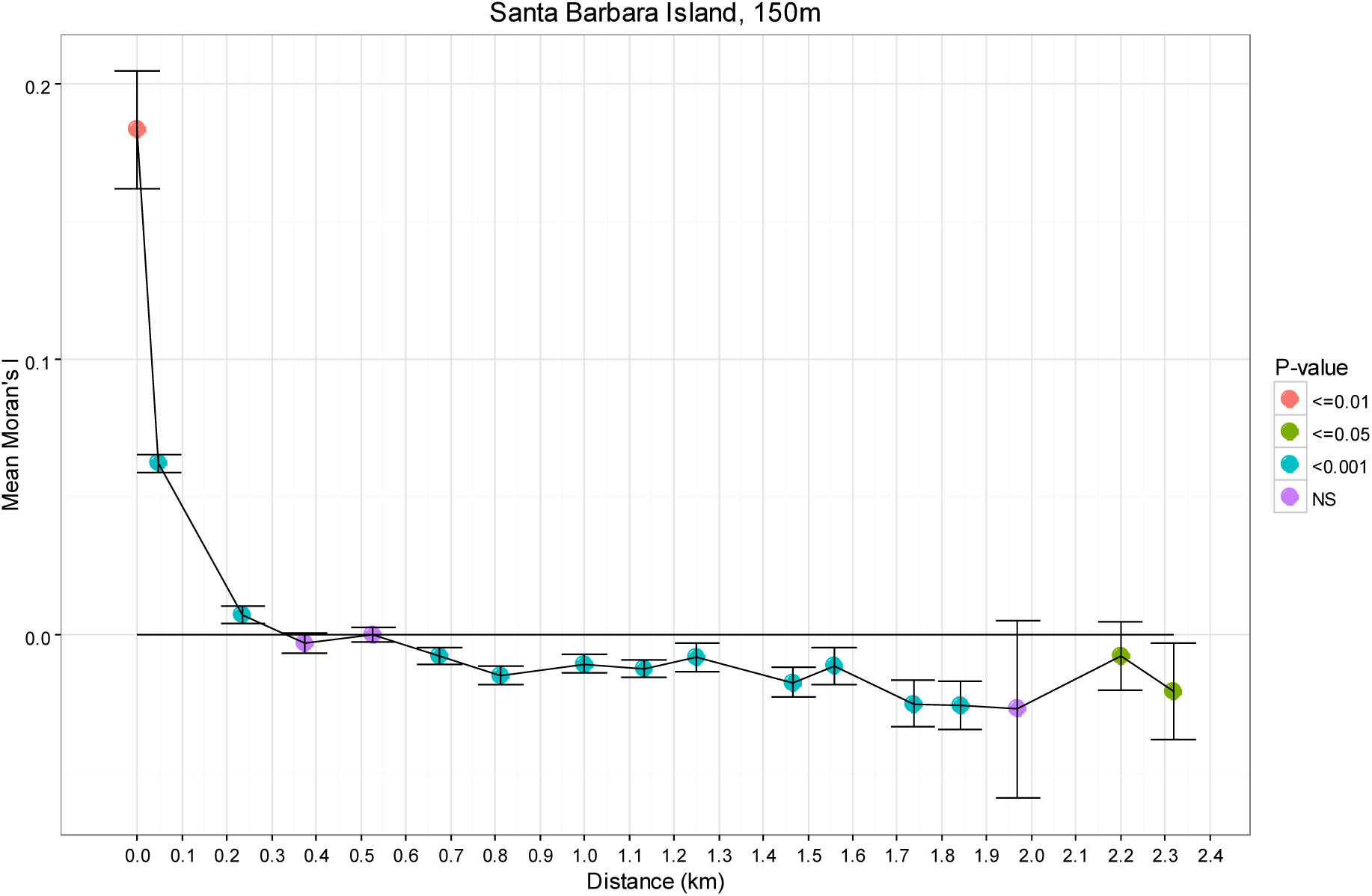
Correlogram of Moran’s I on Santa Barbara Island. Moran’s I was calculated for each distance bin and significance assess through permutation. Points represent the value of Moran’s I for the mean distance within each 150 m distance interval. Bars indicate standard error around the mean while each point color indicates significance of correlation within each point.

**Figure 2:**
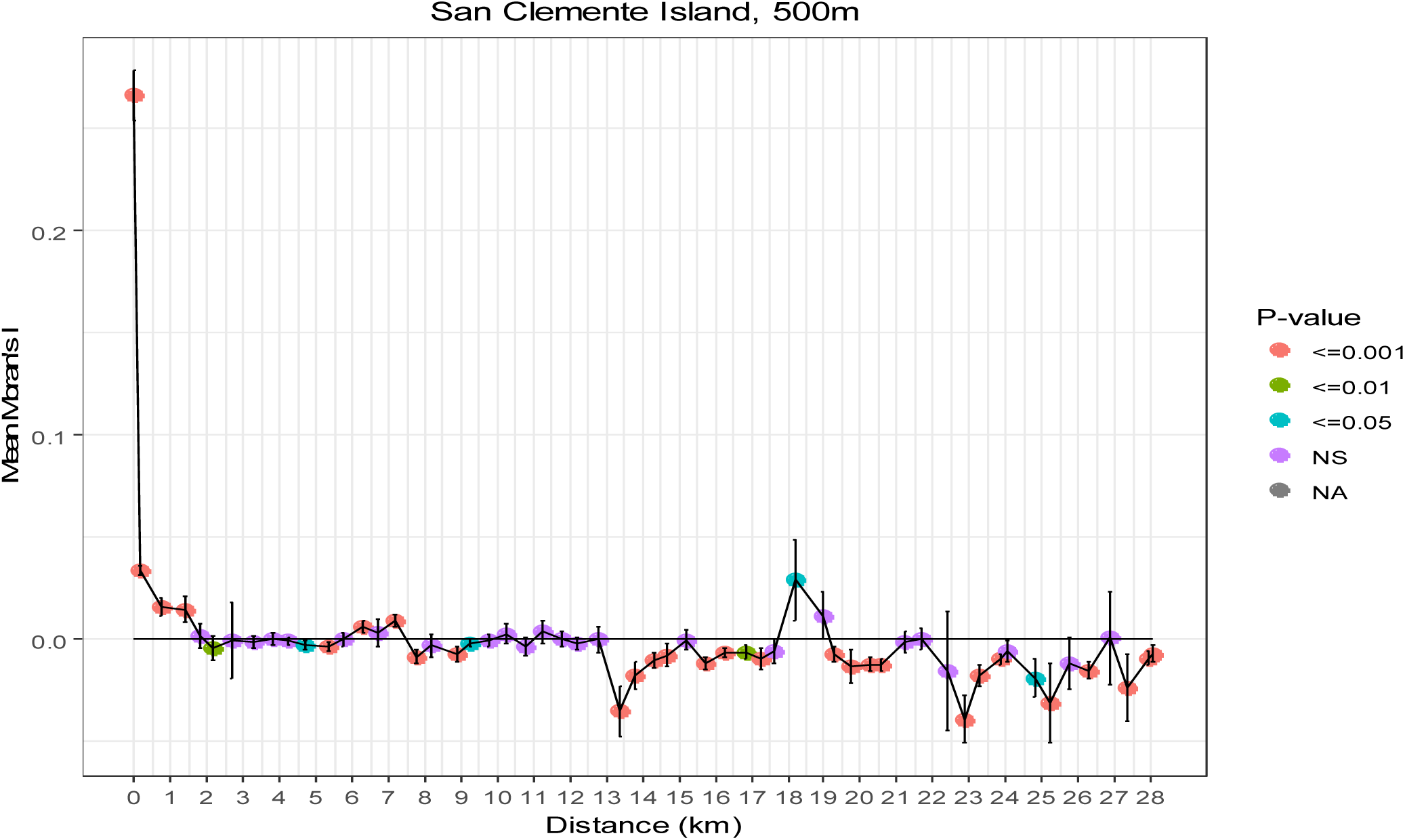
Correlogram of Moran’s I on San Clemente Island. Moran’s I was calculated for each distance bin and significance assess through permutation. Points represent the value of Moran’s I for the mean distance within each 500 m distance interval. Bars indicate standard error around the mean while each point color indicates significance of correlation within each point.

**Figure 3:**
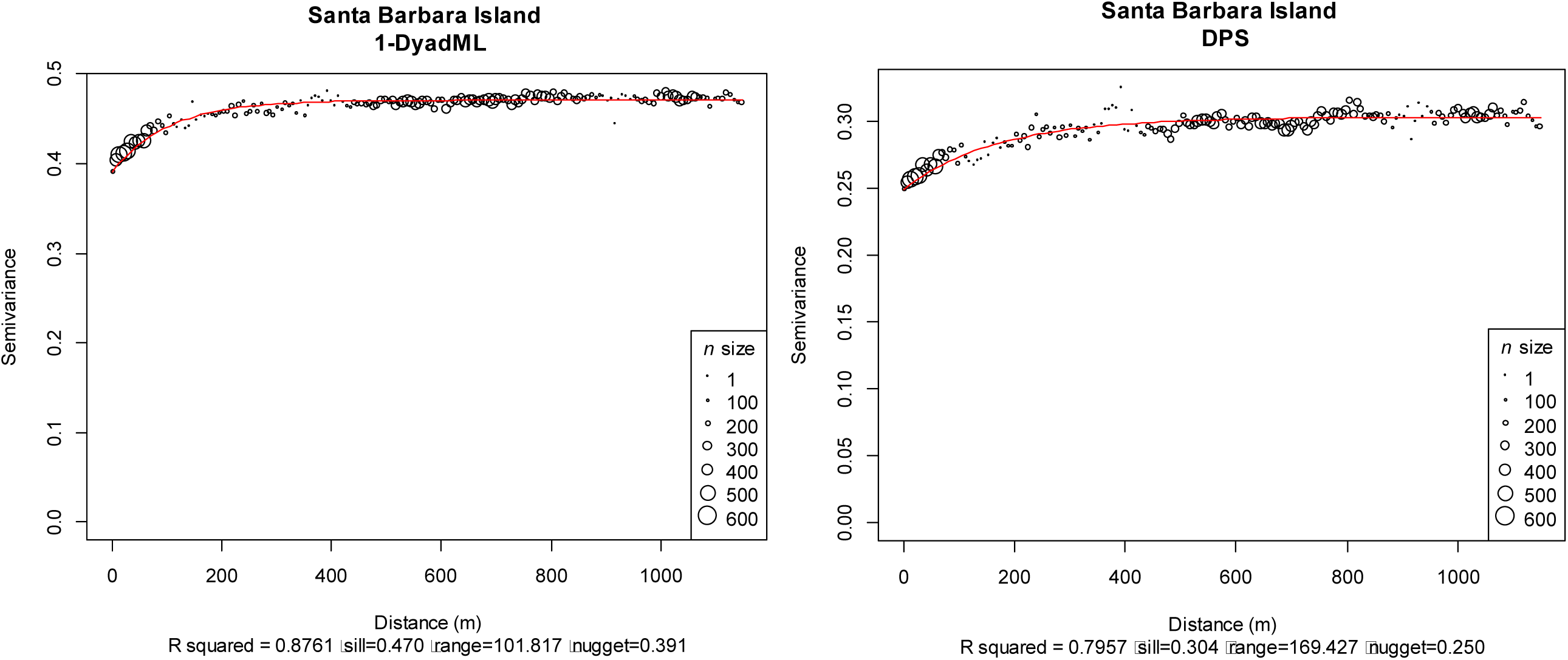
Variogram Analyses for Santa Barbara Island. Empirical variograms (circles) were generated from two distance measures: left) 1-DyadML relatedness estimates, right) Inter-individual genetic distance DPS. Circle size denotes the number of pairwise comparisons within each distance class. Maximum distance evaluated was 1,100 m with a lag distance of 7 m for both variables. Theoretical variograms (red line) were fit as fixed-nugget models. Model R^2^ values were used as a measure of model support and the point of spatial independence is denoted by the range.

**Figure 4:**
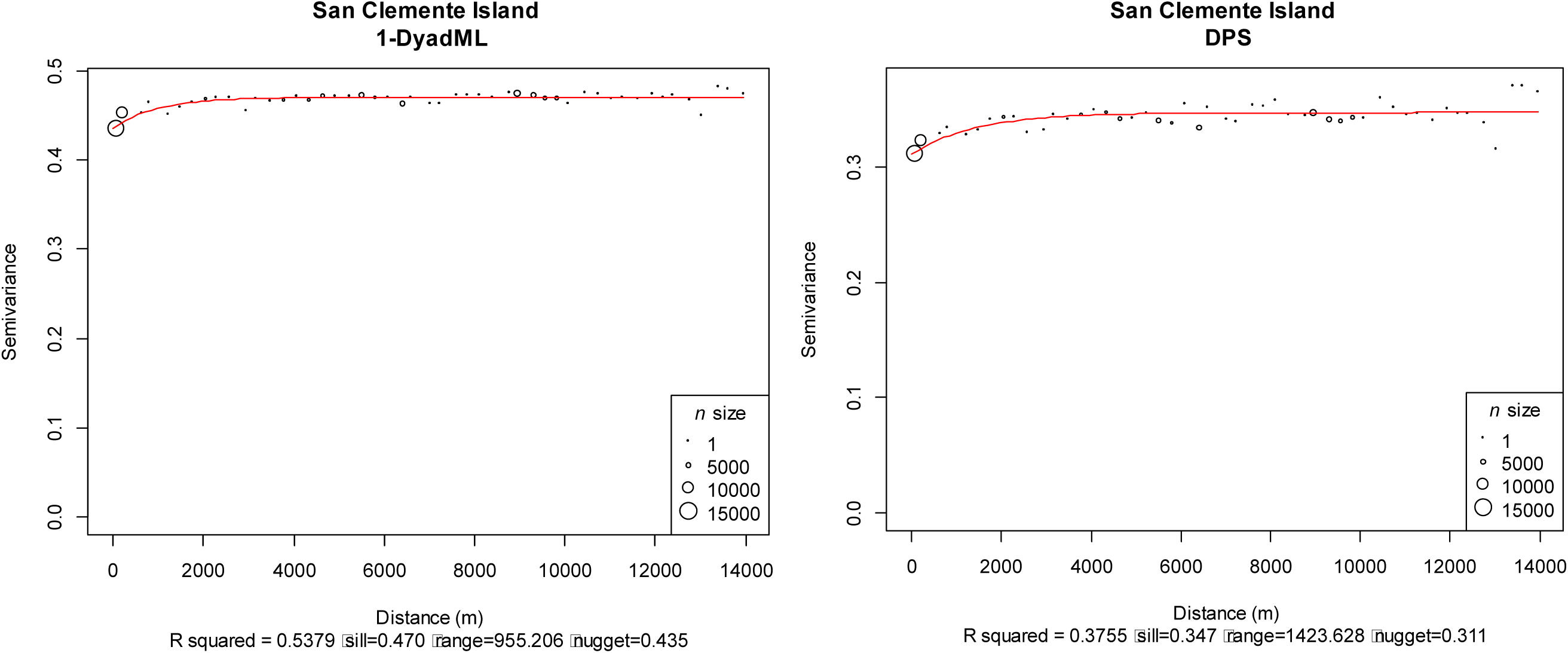
Variogram Analyses for San Clemente Island. Empirical variograms (circles) were generated from two distance measures: left) 1-DyadML relatedness estimates, right) Inter-individual genetic distance DPS. Circle size denotes the number of pairwise comparisons within each distance class. Maximum distance evaluated was 14,000 m with a lag distance of 290 m for both variables. Theoretical variograms (red line) were fit as fixed-nugget models. Model R^2^ values were used as a measure of model support and the point of spatial independence is denoted by the range.

We found no statistical support for sex-biased dispersal on either island. We utilized LRDM in both exploratory and predictive contexts. Exploratory analyses revealed that relatedness and relationship were the only significant predictors of co-capture when controlling for young of year on each island (Table 3). Pairs of individuals were 1.067 times more likely to be captured at the same point with a one standard deviation change in DyadML relatedness estimates on SBI and 1.031 times more likely on SCI. To examine the predictive ability of these patterns we utilized a traditional logistic regression with SBI as the training dataset and SCI as the test dataset. This predictive analysis DyadML and COLONY-inferred relationships had high accuracy (0.9996) and low misclassification (0.0004) rates when predicting capture success trained on SBI and tested on SCI.

**Table 3:** Logistic Regression on Distance Matrices Results. LRDM was conducted separately for each island (SBI=Santa Barbara Island, SCI=San Clemente Island) and for each data set with all individuals (All Samples) or controlled by removal of young of year (>40mm SVL). LRDM was conducted on each predictor variable (Model) independently to determine significance (p-value) through permutation and model support by Nagerkerke’s R^2^ (R^2^). Odds-ratios (Odds-ratio) were computed from 3-variable models in which either the DyadML estimator (Dyad) or COLONY-inferred relationships were used (COLONY) due to the collinear nature of relatedness and relationship. Since the predictors of sex and maturity were used in 2 3-variable models both odds ratios are listed with the forward slash separating the DyadML model from the COLONY relationship model. The predictors of sex and maturity were categorical matches between male and female (sex) and sexually mature and immature individuals (Maturity).

|  | SBI |  |  |  |  |  | SCI |  |  |  |  |  |
| --- | --- | --- | --- | --- | --- | --- | --- | --- | --- | --- | --- | --- |
|  | All Samples |  |  | >40mm SVL |  |  | All Samples |  |  | >40mm SVL |  |  |
| Model | p-value | Odds-ratio | $R^2$ | p-value | Odds-ratio | $R^2$ | p-value | Odds-ratio | $R^2$ | p-value | Odds-ratio | $R^2$ |
| *Dyad* | 0.0001 | 1.0644 | 0.112 | 0.0001 | 1.0659 | 0.112 | 0.0001 | 1.4373 | 0.339 | 0.0001 | 1.0307 | 0.081 |
| *COLONY* | 0.0001 | 1.7004 | 0.089 | 0.0001 | 1.7316 | 0.089 | 0.0001 | 0.9969 | 0.392 | 0.0001 | 0.9990 | 0.073 |
| Sex | 0.6845 | 1.3120/<br>2.4283 | 0.008 | 0.7005 | 1.3265/<br>2.4386 | 0.008 | 0.0001 | 1.0006/<br>1.0003 | 0.018 | 0.3911 | 1.0003/<br>1.0002 | 0.005 |
| Maturity | 0.9356 | 1.3259/<br>1.2964 | 0.001 | 0.8512 | 1.3409/<br>1.3139 | 0.001 | 0.0033 | 1.0005/<br>0.9799 | 0.009 | 0.6900 | 1.0003/<br>0.9959 | 0.001 |

## DISCUSSION

We found inferred parent-offspring distances were between 2 and 13 times greater than the individual displacement distances of long-term CMR studies of 3 to 6 m (Fellers & Drost 1991; Mautz 1993). We provide the first quantitative estimates of dispersal distances in the species (14 m on SBI and 41 m on SCI) and found spatial autocorrelation of allele frequencies, genetic distance, and relatedness at previously unrecorded scales. The LRDM approach of Prunier et al. (2015) resulted in relatedness and relationships being the strongest predictors of island night lizard co-captures, suggesting a social structure with kin-affiliative behavior.

### Distances Between First-Order Relatives

Fellers and Drost (1991) found 12 juveniles on SBI 30 to 40 m from prime habitat over a 6 year study and surmised these represented juvenile dispersal, whereas individual recapture data suggested average displacements of 5.6 m. While there is no indication of juvenile dispersal distances for SCI, Mautz (1993) found that individual relocations to be only an average of 3 m apart, with the longest recorded displacement at 18.5 m. Our findings provide the first quantitative estimates of juvenile dispersal distances, although the distance between parent-offspring pairs is approximately 3 times shorter than the estimates of Fellers and Drost (1991). However, our estimates of dispersal distance are approximately 2.5 times greater than the individual movement distances on SBI and 13 times greater than those on SCI.

Moore et al. (2014) and Dussex et al. (2016) both found parentage analyses more accurate in describing dispersal ecology than alternate genetic methods. Our results support the utility of parentage and kinship analyses in characterizing natal movement, especially when focused on parent-offspring pairs. Estimation of first-order relationships as described in Rice and Clark (2016) and their pairwise differences may be informative in the characterization of dispersal when few parent-offspring comparisons are available. Comparisons of mean pairwise distance for COLONY-inferred relationships were not significantly different between parent-offspring or full sibling groups for either island. The mean distances inferred for first-order relationships differed between islands, potentially due to differences in island scale and population densities. It is notable that the maximum distance between first-order relatives is very close (SBI=158.63 m, SCI=156.75 m) between both islands, although these maximums belong to different relationship groups (full sibling and parent-offspring respectively) for each island.

### Gene Dispersal Distances

Estimates of gene dispersal were also remarkably close between each island at the same lag distances and density estimate methods. Comparing these two metrics, distances from inferred parentage and sigma estimated distances were within 3 m on SBI and SCI when evaluating distances with young of year included, and remained consistent when young of year were excluded. The discrepancy between parent-offspring distance and σ for SCI when controlling for young of year may be attributable to changes in sample size and analytical method. Future research should consider simulation-based approaches to evaluate the accuracy of these metrics compared to the known simulation parameters; however, in the context of estimating parameters for natural populations characterized by an isolation by distance pattern, parentage analyses and σ both appear to yield consistent results.

### Spatial Autocorrelation

Spatial autocorrelation analyses, such as Moran’s I, have been used to understand the scale of autocorrelation for individual allele frequencies and interpret this scale as a measure of dispersal (e.g. Epperson & Li 1997; Yaegashi et al. 2014). The interpretation of the correlogram’s x-intercept as average maximum dispersal distance is uncommon in the literature, and generally noted as the scale at which allele frequencies are become spatially independent. This method returned notable differences between islands, with SBI reaching this point at 375 m and SCI at 1,813 m. These scales are much larger than the displacement estimates from long-term field studies and are also much larger than our dispersal estimates from parentage analyses and σ. Thus, we recommend that studies focused on estimating dispersal distances should favor parentage analyses or gene dispersal distances over spatial autocorrelation analyses.

### Variograms

Estimates using variograms were also incongruent with parentage analyses, but provided smaller estimates of the scale of spatial autocorrelation than Moran’s I. The use of variograms to estimate dispersal has not been formally studied. These estimates could denote the scale of spatial genetic structure (Wagner et al. 2005), connectivity among localized groups (Le Corre et al. 1998), or the “patch size” of the process evaluated (Legendre & Fortin 1989). Because we generated distance measures from the proportion of shared alleles and relatedness our estimates may denote the “patch size” of relatedness, or may indicate familial territories. More meaningful biological interpretation of these field studies using long-term telemetry paired with parentage analyses would provide data on individual movement necessary for more thorough interpretation of genetic patterns. However, simulation-based approaches may offer a more tractable solution to determine whether range estimates produced from spatial autocorrelation and variogram analyses are congruent with known or simulated dispersal patterns.

### LRDM

The results of LRDM indicate that pairwise relatedness and relationships are the best predictors of capturing individuals together. Due to the late spring and summer sampling on SBI the removal of young of year individuals had little effect on the pseudo-R^2^ or odds ratio. However, extensive sampling on SCI during the autumn to capture neonates and associated adults impacted both the pseudo-R^2^ and odds ratio for relatedness, but this remained a significant predictor even after removal of young of year. The continued association of relatives after the ca. 0.8 year mark suggests a level of previously undocumented sociality within the system, which may explain the strong signals of isolation by distance reported by Rice and Clark (2016).

Cryptic sociality and kin affiliation has been noted for the sister species (*X. vagilis*) on the mainland, in which fostering was demonstrated to effect philopatry and kin-affiliative behaviors through delayed dispersal (Davis 2011; Davis et al. 2010). The studies of Davis et al. (2010) highlight several similarities shared between the two species, including dense populations, low dispersal, and small home range sizes. Thus, our findings suggest island night lizards may also form kin groups through delayed juvenile dispersal and prolonged parent-offspring interactions as noted by Davis et al. (2010). Some reclusive reptiles that are social also exhibit parental care, such as attendance of neonates in egg-guarding lizards (e.g. Huang 2006; Mateo & Cuadrado 2012) or maternal attendance of pre-ecdysis neonates in pit-vipers (e.g. Greene et al. 2002; Hoss et al. 2015). Given the frequent and prolonged association between adults and neonates, it is possible *X. riversiana* also exhibits parental care. Although predators on both islands are limited, attending parents could protect neonates from intra-specific aggression. Unsurprisingly, samples collected in autumn most frequently included neonates and associated adults as parturition occurs seasonally (Fellers & Drost 1991; Mautz 1993). However, the association of related island night lizards extending beyond this parturition period warrant further investigation into the extent of their social structure and affiliative behaviors.

### Conservation Implications

Our characterization of dispersal ecology of the island night lizard suggests scale dependent effects and supports the independent management of each island. On SBI, we found parent-offspring distances of approximately 14 m and spatial autocorrelation up to 375 m whereas dispersal distances on SCI were 41 m with spatial autocorrelation up to 1,813 m. These distance estimates will be useful for the management and post-delistment monitoring of populations on both islands (USFWS 2014). We suggest management actions should maintain population sizes and meta-population connectivity, and that the spatial scales derived from our spatial autocorrelation analyses be used to guide those actions. On SBI, a scale of 100 m to 375 m should be used as a focus for remediation efforts, such as direct-line connectivity between habitat patches. On SCI, management should focus at the scales of 1-2 km in efforts to connect remote patches through corridors of prime habitat, as opposed to replanting isolated or remote patches. Furthermore, these findings can inform the design and mitigation of increased infrastructure by identifying patches that would become disconnected at these scales under increased development.

### Conclusions

Dispersal is a key factor in the life history of a species, and a key parameter affecting conservation and management decisions (Bowler & Benton 2005; Hawkes 2009). Although dispersal can be directly observed, it is often labor and time intensive. Indirect inference of dispersal based on genetic evidence is gaining in application but lacks a framework for consistent and reliable inference. Studies utilizing both CMR and genetic inference methods have found that CMR methods generally underestimate dispersal distances, while assignment methods on genetic data can often overestimate dispersal and conflict with direct observations (e.g. Dussex et al. 2016). The application of genetic inference methods to estimate dispersal is likely to be a valuable tool for conservation management in understanding the scale of dispersal processes and the potential effects of management actions on connectivity. However, we have demonstrated that different inference methods may yield very different results which may lead to incorrect inferences and misspecification of parameters, rendering management actions ineffectual (Bowler & Benton 2005; Jackson & Fahrig 2014). Recent studies found parentage analyses to be the most accurate method for characterizing dispersal ecology, and our analyses of the island night lizard support this usage. Furthermore, our findings suggest that natal dispersal parameters should not be derived from spatial autocorrelation or variogram analyses as the parameters inferred are highly variable and likely overestimate dispersal in the context of individual movement. However, these approaches may yield insight into the scale of fine-scale patterns relevant to conservation and suggest a minimum scale below which individuals are likely to be related and thus management actions may be confounded (e.g. Jackson & Fahrig 2014).

## ACKNOWLEDGEMENTS

Support for this research came from United States Department of Defense (Award Number W9126G-12-2-0060) and the Southern California Research and Learning Center (Award Number S18309). We would like to thank the National Park Service for access to SBI, SCI Naval Base for access to SCI and our field assistants. In addition we would like to thank Drs. A. Bohonak (SDSU), H. Regan (UCR), K. Anderson (UCR), and J. Gatesy (UCR) for comments received during manuscript preparation.

## SUPPLEMENTALS: RAW DATA GRAPHS

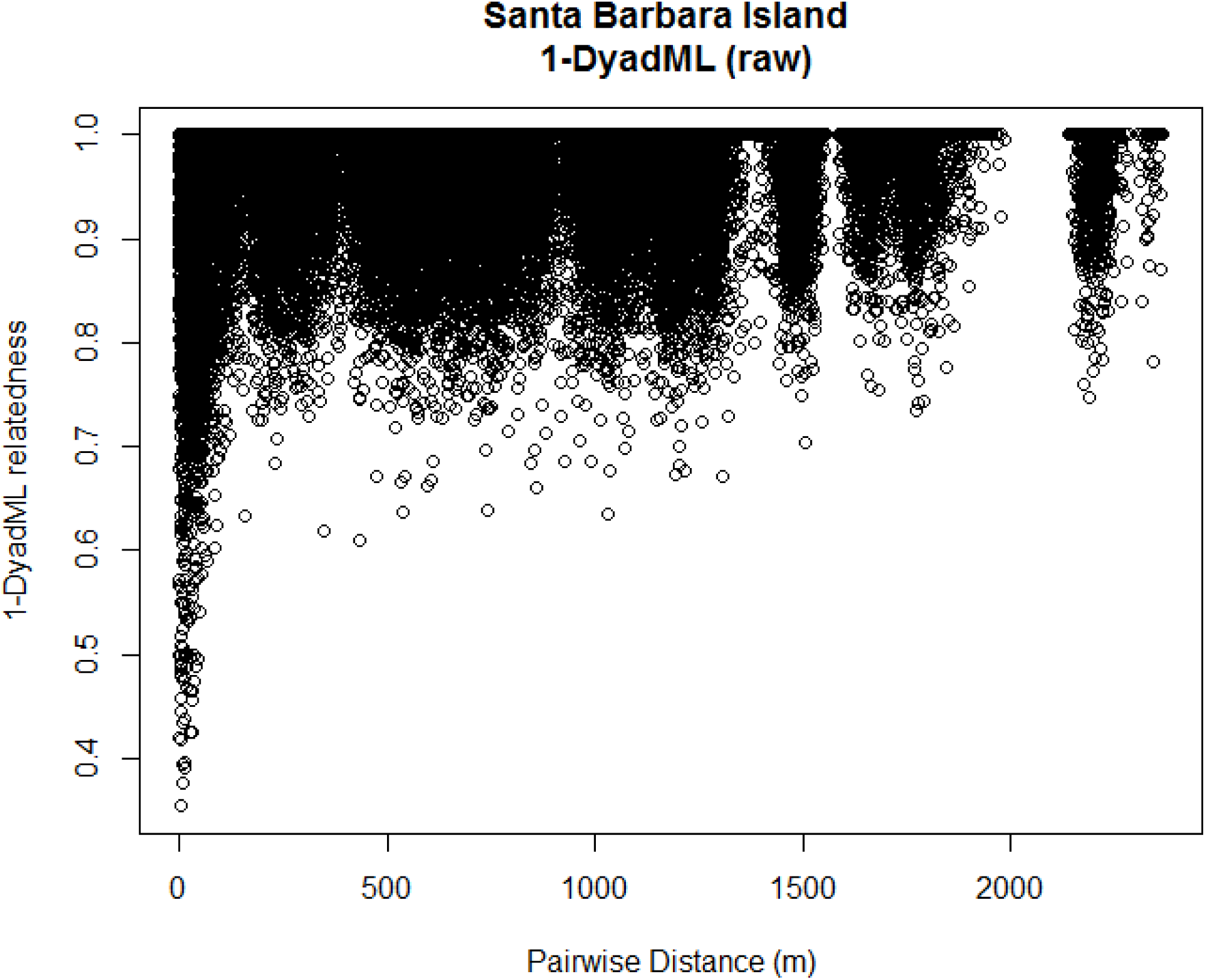

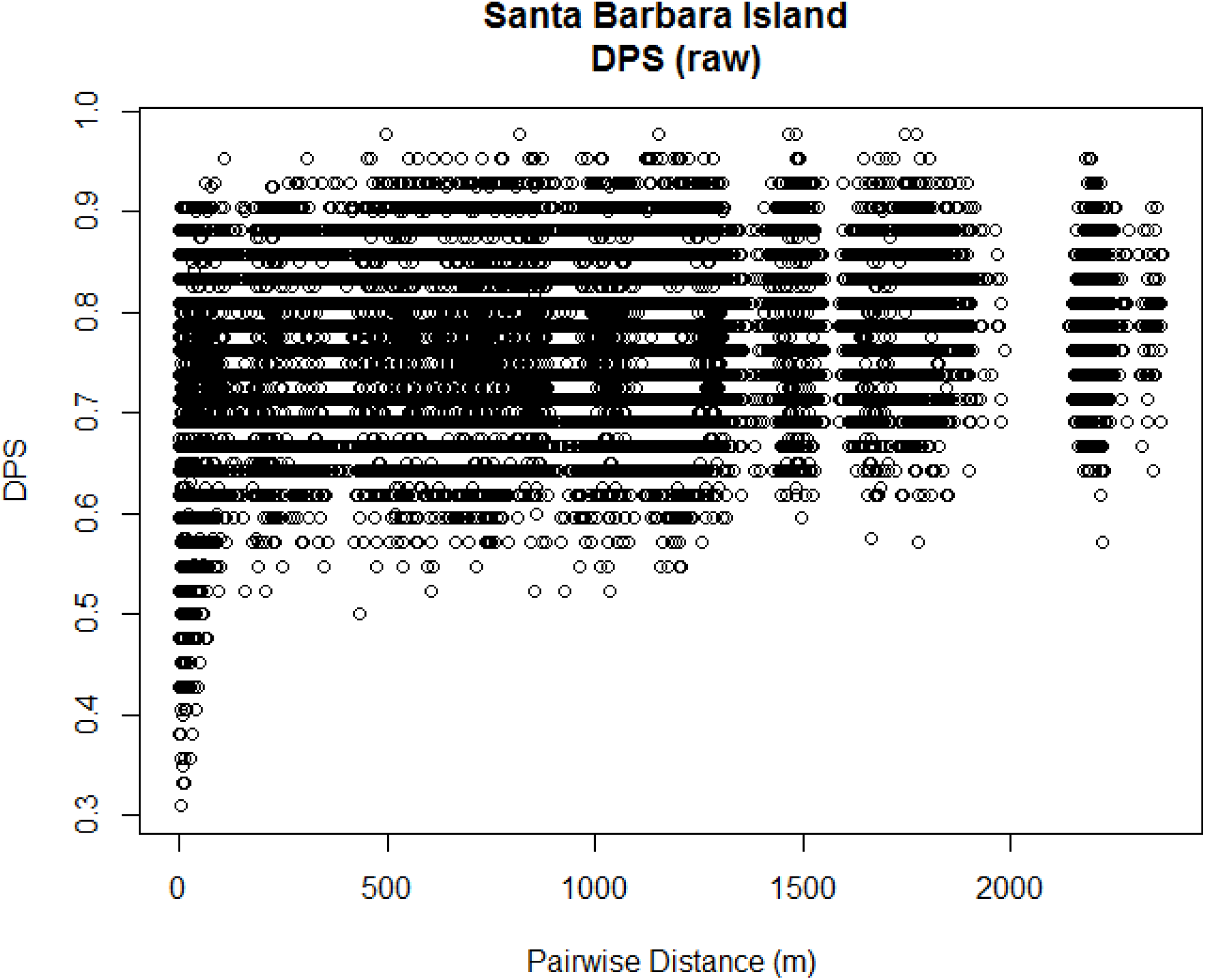

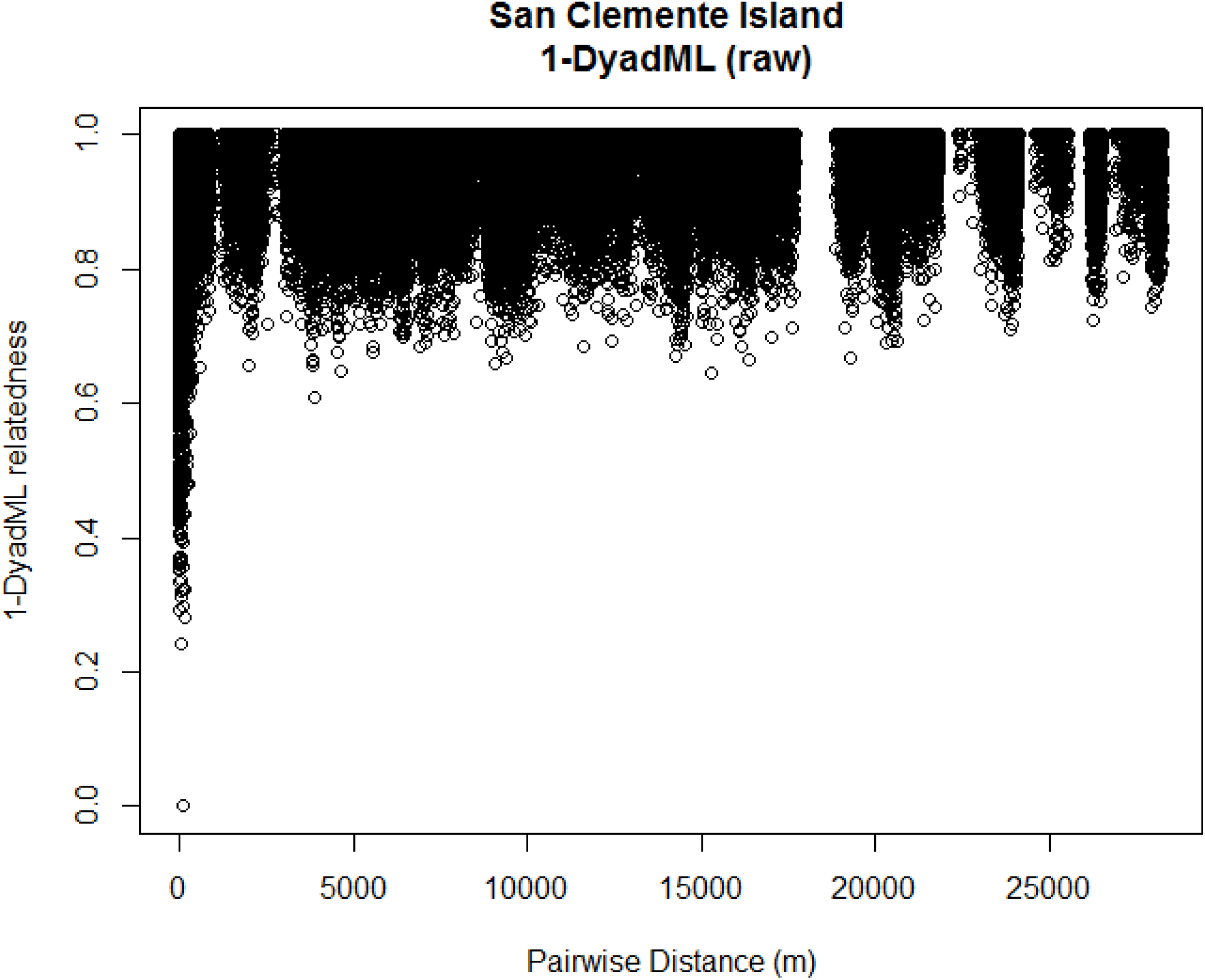

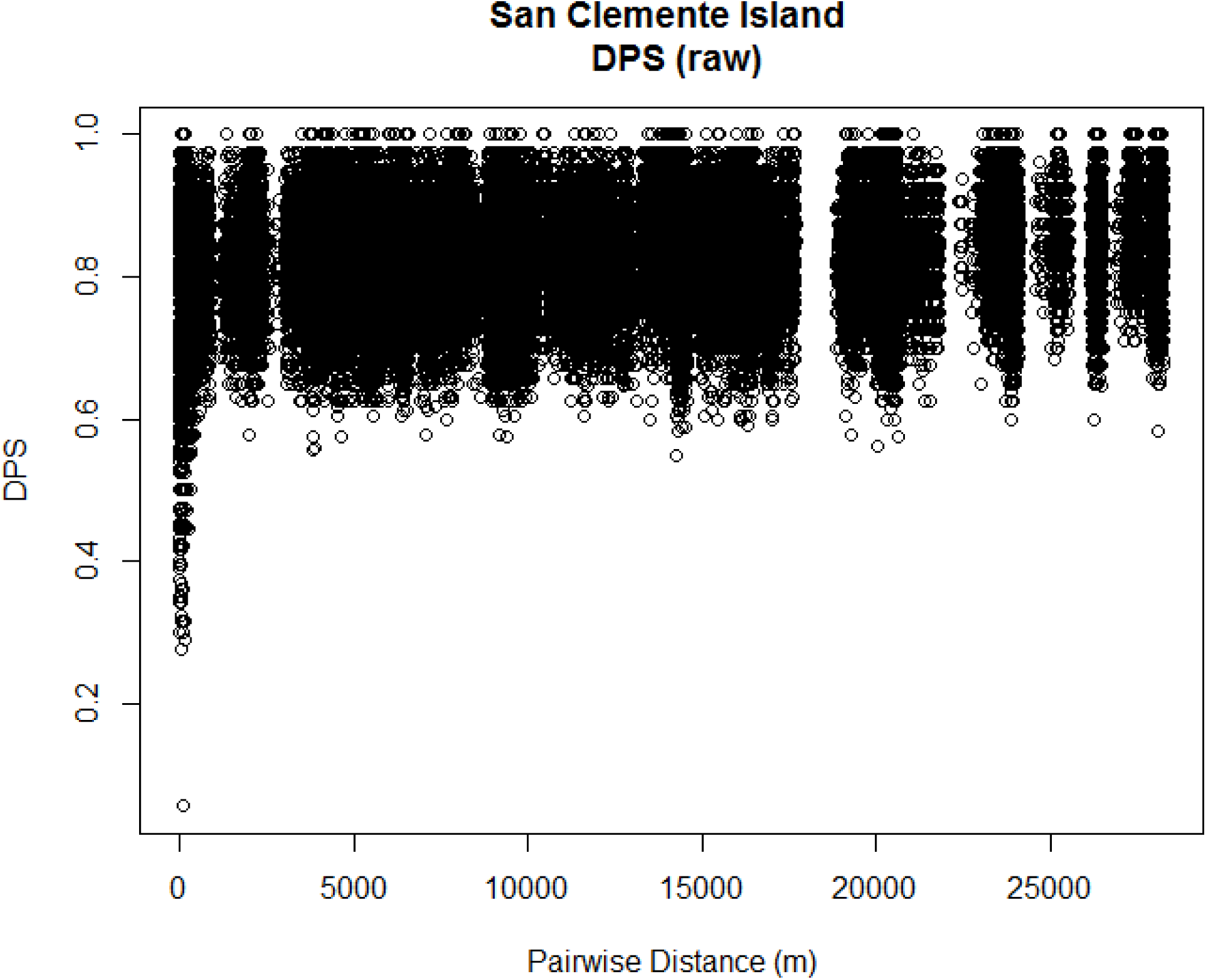

